# Investigation of the spatial heterogeneity of soil microbial biomass carbon and nitrogen under long-term fertilizations in fluvo-aquic soil

**DOI:** 10.1101/493973

**Authors:** YE Hong-ling

## Abstract

Soils are heterogeneous and microbial spatial distribution can clearly indicate the spatial characteristics of the soil carbon and nitrogen cycle. However, it is not clear how long-term fertilization affects the spatial distribution of microbial biomass in fluvo-aquic soil. We collected fluvo-aquic soil samples (topsoil 0-7.5 cm and sub-topsoil 7.5-20 cm) using a spatially-explicit design within three 40.5 m^2^ plots in each of four fertilization treatments. Fertilization treatments were: cropping without fertilizer inputs (CK); chemical nitrogen, phosphorus, and potassium fertilizer (NPK); chemical fertilizer with straw return (NPKS); and chemical fertilizer with animal manure (NPKM). Variables included soil microbial biomass carbon (MBC) and nitrogen (MBN), and MBC/MBN. For both soil layers, we hypothesized that: microbial biomass was lowest in CK but with the largest spatial heterogeneity; and microbial biomass was highest in NPKM and NPKS but with the lowest spatial heterogeneity. Results showed that: (1) Fertilization significantly increased MBC and MBN more in topsoil than sub-topsoil but had no MBC/MBN changes. (2) The coefficient of variation (CV) and Cochran’s C showed that variation was largest in CK in topsoil and NPK in sub-topsoil and that variation of topsoil was generally lower than in sub-topsoil. The sample size of the three variables was largest in CK in topsoil but had little variation among the other treatments. (3) The trend-surface model showed that within-plot heterogeneity varied substantially with fertilization (NPKM = NPK > NPKS > CK), but Moran’s I and the interpolation map showed that spatial variability with fertilization followed the order NPK > NPKS > CK = NPKM at a fine scale in topsoil. In sub-topsoil, the trend-surface model showed that within-plot heterogeneity followed the order NPKM = CK > NPK > NPKS and that the fine-scale pattern was NPKM>NPK=NPKS>CK. MBC had the highest spatial heterogeneity among the three variables in both soil layers. Our results indicate that the application of organic fertilizer (straw or manure) reduced the variation of MBC and MBN but increased the spatial variability of MBC and MBN. The spatial variation of the three variables was MBC > MBN > MBC/MBN regardless of whether variation was considered at the plot-scale or the fine-scale in both layers.

## 1. Introduction

With the increasing application of chemical fertilizers in recent decades, fertilizer efficiency has gradually decreased. This phenomenon is especially common in the North China Plain, which is mainly dominated by fluvo-aquic soil [1, 2]. Some studies have shown that fertilization, especially straw return and organic manure, can effectively improve the fertility of fluvo-aquic soil [3-6]. Fertilization has major effects on soil properties, including improving soil physical structure, increasing soil available nutrients, and increasing soil microbial activity [7-9], and can modify the spatial distribution of soil properties [7, 9, 10].

Soil microorganisms are key factors in the degradation of soil organic matter and nutrient cycling [11, 12]. Soil microbial indicators are very sensitive to the effects of human activities on soil, especially in agricultural activities [13, 14]. With the development of precision agriculture, the spatial distribution of soil microorganisms frequently affects efficient use of nutrients [15, 16] and the spatial heterogeneity of microbes strongly influences the infiltration, migration, and adsorption of soil nutrients [17, 18]. Fluvo-aquic soil is derived from alluvial soil and thus has greater spatial heterogeneity than aeolian soil [19]. Spatial variation in soil can be detected, estimated, and mapped using geostatistical analysis [20], which can help identify changes in spatial trends and characteristics of space in the soil.

Compared with physical and chemical properties, soil microbes are more susceptible and more sensitive to the influence of external sources of nutrients [21]. Soil microorganisms exhibit high spatial and temporal variation, even in homogenously managed agroecosystems [22-24]. At present, studies of spatial heterogeneity of soil microorganisms are mainly conducted in forests [25, 26], grasslands [27, 28], or large scale farmlands [29]. However, even at the small scale level, there is large spatial variability of microorganisms in the soil [30, 31]. Spatial variability of soil properties are controlled by inherent variations in soil characteristics and are affected by exogenous factors such as cultivation of crops or grazing of large herbivores [32-34]. Soil microbial heterogeneity can lead to spatial dependence in soil microbial communities at multiple scales, ranging from the rhizosphere to scales of more than 100 m [35-38]. Previous work has found that variation in microbial biomass can be partially explained by soil type and drainage class [39, 40], soil texture [37, 41], organic matter content, and pH [34, 42]. The conventional plow depth is 20 cm [43, 44], but roots reach from 0-30 cm [45, 46], and microbes are usually present in the topsoil (0-7.5 cm) [47-49]. This spatial pattern leads to variation between the strongest microbial activity and the depth of nutrient uptake by plant roots, so it is necessary to study differences in the spatial distribution of microbes in the topsoil and sub-topsoil layers. The spatial distribution and structure of soil properties can provide us with a better understanding of the potential factors (e.g., grazing, cultivation, conservation, fertilization, and plant density) that influence the other soil properties [35].

For long-term fertilization of farmland, the availability of external materials, especially carbon and nitrogen sources, has become an important guideline of microbial distribution. Röver, et al. [9] focused on changes in spatial heterogeneity of a microbial community due to different fertilization treatments in a small farmland in Germany. They found that banding fertilizer caused microbial biomass carbon (MBC) to reach more than 15%, a moderate degree of variation, forming “hotspots” in some areas. Heinze, et al. [7] showed that organic fertilizer improved the stability of soil pH compared with a single application of chemical fertilizer, however, soil heterogeneity lead to spatial heterogeneity of soil pH, which ultimately masked the effect of fertilization on soil microbial biomass. Wang, et al. [50] found that variation of soil MBC had a strong spatial autocorrelation with a dependence distance of 3.17 m, which was the smallest among all variables (e.g. soil respiration, soil moisture) in 16×14 m plots.

The spatial heterogeneity of soil MBC and microbial biomass nitrogen (MBN) in different fertilization treatments can be used to determine the spatial characteristics of long-term soil evolution. Different fertilization treatments in long-term studies result in uneven distributions of nutrients in the soil, which may further lead to spatial variation of microbial biomass distribution. Dissolved organic carbon and dissolved organic nitrogen vary greatly among treatments, which is the main factor limiting microorganisms[51]. Treatment without fertilizer (CK) has small concentrations of MBC and MBN due to limited sources of carbon (C) and nitrogen (N) and root stubble redistributes the spatial heterogeneity of microbial biomass. Chemical nitrogen, phosphorus and potassium fertilizer (NPK), chemical fertilizer with manure (NPKM), and chemical fertilizer with straw return (NPKS) have different C and N sources. There is sufficient C and N in NPKM in both soil layers, meaning that NPKM may have large microbial biomass and relatively small spatial heterogeneity if fertilization is uniform. But the structure of organic matter in manure is complex and it usually cannot be used directly by microorganisms [52], and on the other hand, simple cellulose and hemicellulose in straw can be decomposed easily [53, 54], so the soil dissolved organic matter may have a larger spatial heterogeneity in NPKM than NPKS. Root stubble is the main carbon source in NPK, and its quantity is very small compared with NPKS and NPKM but it is more than CK, so its spatial heterogeneity should be higher than NPKS and NPKM but lower than CK.

Our objective was to examine the effects of long-term fertilization on spatial heterogeneity of soil MBC, MBN and MBC/MBN in both topsoil and sub-topsoil. In view of previous studies, our hypothesis is that spatial heterogeneity in both layers will be in the following order: CK>NPK>NPKM>NPKS.

## 2. Materials and methods

### 2.1 Site description

Soil samples were taken from the national soil fertility and fertilizer efficiency long-term monitoring station, located in Yuanyang County, Henan Province, latitude 35°00′31″N, longitude 113°41′25″E. This site has a mean annual temperature of 14°C; annual precipitation and annual evaporation of 645 mm and 1450 mm, respectively; groundwater depth of 50-80 cm during the rainy season and 150-200 cm during the dry season; and a frost free period of 224 d. In 1991, after two years of uniform planting (1989-1990), the initial soil physicochemical properties were: clay mineral types, hydromica 1.24 g/kg, total porosity 43%; soil texture, silt loam, clay content (< 2 microns) 13.4%, silt content (2-50 microns) 60.7%, and sand content (50-2000 microns), 26.5%; soil pH, 8.3; soil organic carbon 6.7 g/kg; total nitrogen 0.67 g/kg; and soil C/N ratio, 10.

Fertilization treatments included chemical fertilizer nitrogen (N), phosphorus (P), and potassium (K) (NPK); NPK with straw return (NPKS, 70% N from straw before 2004, 50% N from straw after 2004); animal manure and chemical fertilizer NPK (NPKM); and no fertilizer (CK). The cropping system was wheat-maize rotation in one year. Manure and straw were only used in the wheat season. In the wheat season, the fertilization amounts were N-165 kg • hm^-2^, P_2_O_5_-82.5 kg • hm^-2^, K_2_O- 82.5 kg • hm^-2^, with a ratio of N: P_2_O_5_: K_2_O=1: 0.5: 0.5; topdressing occurred at the wheat turning green stage. In the maize season, fertilization amounts were N 188 kg • hm^-2^, P_2_O_5_ 94 kg • hm^-2^, K_2_O 94 kg hm^-2^; with a ratio of N: P_2_O_5_: K_2_O=1: 0.5: 0.5, topdressing was conducted during the grain-filling period.

The topdressing direction went from north to south; the row spacing was 40 cm in the wheat season and 60 cm in the maize season; and the plow depth was 20 cm.

### 2.2 Field sampling and laboratory analysis

There were three replicates in each fertilization treatment, referred to as: CK1, CK2, and CK3; NPK1, NPK2, and NPK3; NPKS1, NPKS2, and NPKS3; and NPKM1, NPKM2, and NPKM3. We studied four fertilization treatments with a total of twelve plots. Fertilization plots were rectangular: the length (south-north) was 9 m and the width (east-west) was 4.5 m. We set the coordinate origin at the northeast corner of every plot; the Y-axis was south and the X-axis was west. We divided every plot into eight squares, with side lengths of 2.25 m. At the center of each square we set a circle with a radius of 1.125 m, and then randomly selected three points within one circle and recorded the coordinate point. These were our sampling points for topsoil and sub-topsoil; the topsoil was 0-7.5 cm underground and the sub-topsoil layer from 7.5-20 cm. In total, 24 samples were collected per plot, 72 samples per fertilization treatment, 288 samples in one layer, and 576 samples in total.

Soil samples were collected on July 11, 2016 (during the maize season). Plots were divided into 8 parts using a measuring rope, and then we positioned cards at sampling locations in the grid. Samples were taken from both soil levels (0-7.5 cm and 7.5-20 cm) in each location using a soil drill. Fresh soil samples were stored in the refrigerator at -20°C. After selecting the debris and sieving using a 2 mm screen, soil MBC and MBN were determined using the chloroform fumigation extraction method. The principle of the method and detailed operation steps are in Vance, et al. [55] and Lin, et al. [56]. Carbon and nitrogen were measured in leachate with a C/N analyzer (multi N/C 3100, Analytik Jena AG, Germany). The formulas used were from Vance *et al.* (1987a):

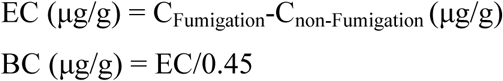

### 2.3 Data analysis

#### 2.3.1 Conventional statistical methods for different fertilization treatments

Means, variances, and frequency were estimated for each soil property in each plot. The distributions of values from three plots in the same treatment were plotted on a common scale for comparison purposes. Cochran’s C test was used to test the assumption of variance homogeneity. The test statistic is a ratio that relates the largest empirical variance of a particular treatment to the sum of the variances of the remaining treatments. The theoretical distribution with the corresponding critical values can be specified [57-59]. The purpose of frequency distribution analysis is to illustrate the overall distribution of soil attribute values in different intervals. Thus, the distribution of MBC and MBN under different fertilization treatments was also analyzed.

#### 2.3.2 Geostatistics methods

We performed spatial statistics using the trend-surface model, Moran’s I, and an interpolation map.

##### 2.3.2.1 Trend-surface model

The trend-surface model is usually used to analyze the spatial rules of soil properties at the plot scale, and then indicates changes in properties on an X, Y coordinate system. The two trend-surface model is more suitable for small scale spatial data analysis. The two trend-surface model is as follows:

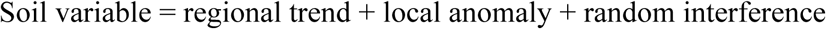

This model sums up the spatial distribution of attributes in order to filter out random interference [60]. According to the complexity of the elements to be evaluated, the model can be divided into three parts: the one trend-surface model, the two trend-surface model, and the three trend-surface model. The study area was small, so the two trend-surface model was used [61]:

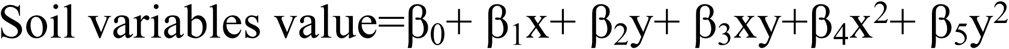

The coefficients of the trend-surface model (β_0_-β_5_) can be obtained by regression: β_0_ is the intercept; β_1_ and β_2_ are the effect of soil property changes in the X and Y axes; β_3_ is the changes in the attribute values on the diagonal; and β_4_ and β_5_ represent the abnormality along the X and Y axes of the parameters to determine the degree of influence of each attribute in each direction of the coordinates. The regression coefficient of the trend-surface model was calculated in SPSS (version 20.0, IBM) at the p<0.05 level.

##### 2.3.2.2 Moran’s I

Moran’s I is the degree of spatial autocorrelation within a plot. Spatial autocorrelation refers to how data are related in space, and whether the relationship of such data is concentrated, dispersed or has no correlation. Spatial autocorrelation analysis is widely used in the field of geographical statistics and the most important indices are Moran’s I and Geary’s C. Moran’s I, which uses the local spatial autocorrelation distance as the maximum variation from different treatments, was the selected. Because the correlation coefficients are between -1 and 1, they are related to the relationship between attribute values within a plot, which is not relevant when the correlation coefficient is 0. Local spatial autocorrelation allows for testing of spatial variability at the small scale [62].

Spatial autocorrelation of Moran’ S I can be expressed as:

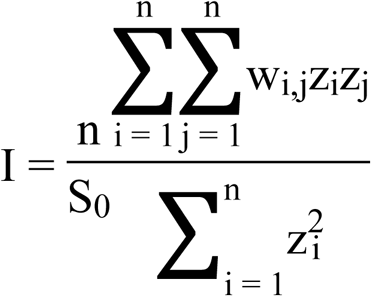

where z_i_ is the deviation of I and its average value 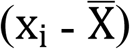, W_i_; y is the spatial weight between i and j; n represents the total number of elements; and S_0_ is the aggregation of all spatial weights.

##### 2.3.2.3 Interpolation map

Interpolation maps are made by converting known discrete measured data to a continuous data surface and then interpolating into the overall plot. The advantages and disadvantages of various interpolation methods are then compared. The Inverse Distance Weighted (IDW) interpolation method does not need the semi variance as a function parameter and the interpolation is inversely proportional to the distance. Therefore, IDW is suitable in small plots of few samples and has high precision [63]; ArcGIS 10.2.1 was used to generate the interpolation map.

## 3. Results

### 3.1 Descriptive statistics

#### 3.1.1 Mean and coefficient of variation

For topsoil, concentrations of MBC and MBN were in the following order: CK < NPK < NPKS = NPKM; there were no significant differences in MBC/MBN among the fertilization treatments (P<0.05). For sub-topsoil, MBC was the highest in NPKM and the lowest in CK. For MBN, there were no significant differences among NPK, NPKS, and NPKM, but it was the lowest in CK. MBC/MBN was highest in CK and lowest in NPK(P<0.05).

The concentrations of MBC and MBN in topsoil were generally higher than in sub-topsoil (P<0.05) (Fig. 2). The MBC concentration was 52.77-67.24% of topsoil in sub-topsoil; MBC concentration decreased fastest in NPKS and slowest in NPK. MBN concentration was 51.71 - 70.95% of topsoil in sub-topsoil(P<0.05). MBC/MBN was only significantly different in NPKM between topsoil and sub-topsoil, where MBC/MBN was 10% lower in topsoil than sub-topsoil(P<0.05).

The coefficient of variation (CV) was typically lower in topsoil than sub-topsoil in all four fertilization treatments (Table 1), except for MBC and MBN of CK. In topsoil, CVs of MBC, MBN, and MBC/MBN were the highest in CK; CV of MBC was the lowest in NPKM; and MBN and MBC/MBN were the lowest in NPKS. In sub-topsoil, the CVs of MBC, MBN, and MBC/MBN were the highest in NPKS, NPK, and CK, respectively, while the CVs of MBC and MBN were lowest in NPKM, and MBC/MBN was the lowest in NPKS.

#### 3.1.2 Frequency

To introduce the effects of fertilization history on soil heterogeneity, individual samples from the three plots of each fertilization treatment were pooled (n=72), illustrating the frequency distributions of soil properties (Fig. 3).

The frequency of MBC in topsoil was the highest in 200-300 mg/kg for CK, between 300-500 mg/kg for NPKS and NPKM, and between 200-400 mg/kg for NPK. MBC in sub-topsoil was the highest in 100-200 mg/kg for CK, 200-300 mg/kg for NPKM and NPKS, and 100-300 mg/kg for NPK.

The frequency of MBN in topsoil was highest in 20-60 mg/kg for CK, and between 40-80 mg/kg for NPK, NPKS, and NPKM. MBN in sub-topsoil was the highest in 0-40 mg/kg in CK, 20-40 mg/kg for NPK, and NPKS and NPKM had the highest frequency between 20-60 mg/kg.

For MBC/MBN, these four treatments were concentrated at 4-12 in topsoil and sub-topsoil.

In Fig. 3, we can also see that MBC and MBN are more concentrated in sub-topsoil than topsoil, and the distribution range of MBC was relatively narrow due to its low concentration in CK. The distribution of MBC in NPKM was more concentrated than in NPKS. The highest concentration of MBC in sub-topsoil still occurred in NPKM, and the lowest concentration was in CK.

#### 3.1.3 Within-plot variances

Coefficients of variation (CVs) of soil properties within-plot are summarized in Fig. 4. The CVs of topsoil were mainly in the range of 18-36%, slightly higher in CK and lower in NPKS. CVs were higher in sub-topsoil than topsoil, especially for NPK where three plots were abnormally higher. Overall, MBC/MBN had the lowest CV.

Cochran’s C was used to test the within-plot variance among different treatments or different plots (Table 2). However, there were no significant differences in the variance of MBC in topsoil. The differences in variance of MBN and MBC/MBN were significant. Four of the highest six values of the three variables occurred in CK in both layers, and two of the three maximum values of each variable occurred in CK in topsoil. There were significant differences in variance of the three variables among 12 plots in sub-topsoil. However, we could not confirm which fertilization had the largest variance because the largest variances of the three variables were distributed among different fertilization treatments. For MBN, none of the largest three CV occurred in CK in sub-topsoil, showing higher variability of CK in four fertilization treatments. MBC and MBN were different in sub-topsoil than in topsoil, where they were relatively larger in NPK than the others. MBC/MBN still had higher variation in CK compared with the others.

The Cochran’s C test was also used to compare pairs of treatments conforming to our hypothesis as follows: CK vs. NPKS, CK vs. NPK, CK vs. NPKM, NPKS vs. NPKM, NPK vs. NPKM, and NPK vs. NPKS. The results showed that the three variables of topsoil had higher variation in CK, while MBC had the highest variation in NPKS in sub-topsoil, and MBN had the highest variation in NPK in sub-topsoil, followed by NPKM, NPKS, and CK.

#### 3.1.4 Sample size requirements based on observed within-plot variances

Based on the estimated standard deviation, sample size requirements are different for different fertilization treatments in 40.5 m^2^ plot [64, 65].

In topsoil, the number of samples required for each fertilizer treatment was significantly different (Fig. 5). For MBC, MBN, and MBC/MBN, the sampling size of CK was the largest. When the relative expectation error was 5%, the number of samples required for CK treatment reached 136-424 and the minimum number of samples of NPKS was 88-90. For MBN, with a relative error of 5%, the sample size reached 248-601 in CK but was 128 -132 in NPKS. The sample size of MBC/MBN was lowest relative to MBC and MBN, the maximum sample size of CK3 was 260 and sample size was lowest (48-66) in NPKS when the relative error was 5%.

The sample size was greater in sub-topsoil than topsoil. For MBC, the sample size of NPK reached the maximum (207-561) when relative expected error was 5%, and the sample size was 43-267 at the same relative expected error in NPKM. For MBN, the largest sample size was in NPK at 300-650 with a relative expected error of 5%. The sample size was lowest (between 207-561) in NPKM. MBC/MBN sample sizes did not have large differences; sample size was maximum (382-1016) in NPKS and minimum (81-301) in NPKM.

Sample size requirements depended on long term fertilization (Fig. 5), with more samples required in CK than in NPKS and NPKM in topsoil; NPK and NPKS required the most samples, and NPKM required the fewest, in sub-topsoil.

### 3.2 Spatial statistical analysis

#### 3.2.1 Trend-surface model

The trend-surface model revealed a number of significant patterns in the spatial variability of soil properties among different long-term fertilization treatments.

There was a total of 30 linear or nonlinear relationships in topsoil (Table 3), including ten in NPKM, ten in NPK, nine in NPKS, and one in CK. MBC had a total of 12, MBN had ten, and MBC/MBN had eight. These results indicated that the highest spatial heterogeneity occurred within plots in NPKM and CK had the lowest spatial heterogeneity. MBC had greater spatial variability compared with MBN and MBC/MBN.

Sub-topsoil had greater spatial heterogeneity than topsoil. There was a total of 34 linear and nonlinear relationships in sub-topsoil, CK accounted for nine, NPK for eight, NPKS for eight, NPKM for nine, MBC for 12, MBN for 11, and MBC/MBN for 11. NPKM was the maximum both in topsoil and sub-topsoil and as MBC, MBC/MBN was the minimum both topsoil and sub-topsoil.

#### 3.2.2 Moran’s I

We used local Moran’s I analysis to detect fine-scale spatial structure within each plot after removing coarse-scale trends across the 40.5m^2^ plots. Results are summarized in Table 4.

Topsoil: because there were three replicates and three variables per fertilization treatment, nine plots may have spatial autocorrelation for MBC, MBN, and MBC/MBN in each fertilization treatment. There were three, eight, six, and three plots in CK, NPK, NPKS, and NPKM, respectively, with spatial autocorrelation. MBC, MBN, and MBC/MBN showed stronger spatial autocorrelation in NPK and NPKS. In particular, MBC showed spatial autocorrelation in three plots in NPKS with a spatial lag distance between 1-5.5 m, while only one plot had spatial autocorrelation in NPKM with lag distance between -5.5 and 6.5 m. For MBN, only one plot had spatial autocorrelation in NPKM, but three plots in NPK had spatial autocorrelation, with spatial lag distance between -1-6 m. For MBC/MBN, there was no spatial autocorrelation in CK, and only one plot of three had spatial autocorrelation.

Sub-topsoil: only four of nine plots in CK achieved spatial autocorrelation and there were five, five, and seven in NPK, NPKS, and NPKM, respectively. This suggests stronger spatial autocorrelation in NPKM, especially for MBC, which showed spatial autocorrelation in all three plots, with spatial autocorrelation distances between 4.5-7.5 m. MBN had two of three plots with spatial autocorrelation in CK, NPKS, and NPKM. However, only one plot of three had spatial autocorrelation in NPK. MBC/MBN was similar to MBN: each fertilization treatment had two of three plots with spatial autocorrelation in NPK, NPKS, and NPKM, while CK only had one of three.

#### 3.2.3 Interpolation map

We used two-dimensional interpolation maps to visualize MBC, MBN, and MBC/MBN by inverse distance weighting (IDW) to better estimate soil in the whole plot. This analysis was done to compare the spatial distributions of the soil properties among the plots.

The interpolation maps had darker colors in CK compared to NPK, NPKS, and NPKM in both soil layers and concentrations of MBC and MBN were higher in topsoil than sub-topsoil. The color was more uniform and with few hotspots in CK while there were more hotspots in NPKS and NPKM. The largest hotspots appeared in CK1 in topsoil, with the largest hotspots of MBC and MBN at the bottom right corner of this plot. The largest hotspot of sub-topsoil occurred in NPKM3.

We did not find a significant impact on soil microbial biomass based on the direction of topdressing according our IDW maps, but hotspots were always at the edge of the plots.

## 4. Discussion

### 4.1 Effects of fertilization on MBC, MBN, and MBC/MBN and their variation

MBC and MBN concentration were the highest in NPKM and lowest in CK in both layers, following the order: NPKM = NPKS > NPK > CK. This result was the same as Liu, et al. [66], and MBC and MBN concentration were higher in topsoil than sub-topsoil. There was no significant difference of MBC/MBN among fertilization treatments or between layers (except it was higher in sub-topsoil than topsoil in NPKM). As we know, a large amount of microbial input comes from manure [53, 67] and sufficient dissolved organic carbon and nitrogen caused a high microbial biomass in NPKM. There was also a large amount of microbial biomass in NPKS due to straw providing a sufficient source of carbon and nitrogen for microorganisms. Furthermore, the weakly alkaline soil can neutralize the acids produced by microbes when they decompose the straw [68, 69], producing a soil environment suitable for microorganism growth and propagation and leading to higher MBC and MBN. NPK lacks necessary carbon sources and leads to insufficient energy to supply more microorganisms, and thus has lower microbial biomass. CK had the lowest microbial biomass due to its low carbon and nutrient input. Even in the same tillage layer, topsoil was closer to the ground, with high porosity and good air permeability, which provided better aeration conditions for microbial activity [70, 71]. With increasing depth, the permeability is reduced, which decreases the microbial biomass [72, 73]. The difference of MBC/MBN between the soil layers was not significant except with NPKM. This indicated that the microbial population structure did not change obviously up to a depth of 20 cm. MBC/MBN of topsoil was lower than sub-topsoil in NPKM, indicating that the proportion of fungi increased from topsoil to sub-topsoil, which is likely to be caused by the difference in aeration conditions and available organic carbon in the soil, and manure caused this difference [74-76].

Long-term fertilization treatments showed the highest variation in CK, while NPKS and NPKM had the lowest variation in general, and following the order: CK > NPK > NPKM > NPKS in topsoil and NPK > CK > NPKS > NPKM in sub-topsoil. Variation in sub-topsoil was higher than topsoil except for MBC and MBN in CK. The differences of MBC and MBN among different fertilization treatments matched our hypothesis, probably because of smaller bulk density, macro-water-stable aggregates (>2 mm), high porosity, and because soil moisture and oxygen content were more even in NPKM and NPKS [77-80]. On the other hand, low porosity and low oxygen content were not conducive to microbial survival in CK. A small portion of the soil had high porosity and oxygen content due to root penetration [77, 81] and root stubble, leading microbes to gather near the roots and resulting in an uneven distribution of microbes in the soil. Lateral root systems were developed in NPKM [82, 83], root stubble residues in soil were more balanced, and the overall spatial variability decreased.

### 4.2 Effects of fertilization on the spatial variation of MBC, MBN, and MBC/MBN

Trend-surface models usually indicate spatial variability at the plot scale. In this study, the trend-surface model showed maximum spatial variability in NPKM in both layers and the minimum spatial variability in CK in topsoil. MBC had higher spatial variation than MBN at the plot scale; the minimum was found in MBC/MBN. This means that carbon or nitrogen sources might be not uniform at the plot scale. Manure caused the uneven distribution of dissolved organic carbon and nitrogen in NPKM because manure is difficult to decompose due to being a macromolecule and having high lignin content [84, 85], leading to higher spatial variability at the plot scale. Another reason may be that manure was usually scattered artificially in the plot, which strongly affected the spatial distribution in the plot. The trend-surface spatial properties of CK were almost the opposite in topsoil (smallest) and sub-topsoil (highest), which indicated that as the sole source of soil organic carbon and nitrogen, the distribution of root stubble in the two layers was completely different. The stubble was mainly in sub-topsoil and was relatively less common in topsoil in CK. Spatial variation of MBC/MBN was the lowest, indicating that the microbial communities had no large differences in all soils at the plot scale.

Moran’s I spatial autocorrelation distance represents spatial variability at the fine-scale. There were large differences in spatial heterogeneity between the two soil layers at the fine scale in this study. The spatial variability of NPK was highest, followed by NPKS, CK, and NPKM in topsoil; the spatial variation in sub-topsoil followed the order: NPKM > NPKS = NPK > CK. The only carbon input in NPK was maize root stubble, which is difficult to decompose due its high ligin[86], but microbes tended to gather near the root stubble, resulting in larger spatial variability. Straw was scattered and plowed into the soil in the plot and while the distribution was more uniform in NPKS than NPK, microbial decomposition of straw is uneven: microbial biomass was low in places where straw had higher lignin content or was uncovered, and other places had higher microbial concentration, so the spatial autocorrelation was high at the fine-scale [87, 88]. In sub-topsoil, the strongest spatial autocorrelation occurred in NPKM, possibly due to higher permeability from more dissolved organic carbon and nitrogen [89], resulting in the highest dissolved organic matter content in sub-topsoil. But this phenomenon was inhomogeneous, which is beneficial to the survival and aggregation of microorganisms in certain parts of the sub-topsoil [90, 91]. Straw provides a normal source of energy for microbial survival in NPKS [92-94], but it is more difficult for microbes to survive and the decomposition rate of straw is slower in sub-topsoil [47], soil microbes and other soil organisms also release a variety of organic matter in the soil [95, 96], supplying soil microbial nutrient sources and increasing spatial heterogeneity simultaneously at the fine scale. The root stubble depth also differs among fertilization treatments in sub-topsoil [97-100]. In this study NPKM and NPKS both had high density roots in both layers, but NPK had the same root density in topsoil [101] and the root density was much lower in sub-topsoil. Therefore, the spatial heterogeneity was much lower in NPK. The root density was higher in sub-topsoil than topsoil in CK due to its poor nutrient soil, the root penetrated deeper can it get enough nutrition and moisture[102], which led to very low spatial heterogeneity at the fine-scale in CK in topsoil.

There were different spatial heterogeneity patterns among the treatments at the fine-scale and the plot-scale, so we made a two-dimensional map with the IDW interpolation method to express the spatial distribution of variables in plots. The largest hotspot was in CK1 (high MBC and MBN). This hotspot was caused by residual stubble because we found carbonized maize stubble when picking roots from the soil. In such samples, MBC and MBN were higher, indicating the emergence of microbial aggregation in this position; microbial decomposition was more intense and caused this hotspot. Another obvious hotspot was in NPKS1, but we did not find any abnormal phenomena when we were selecting and sieving. This hotspot might have resulted from rich carbon and nitrogen sources in this sample that had been consumed by microbial decomposition. A notable phenomenon in IDW maps was that the most hotspots were commonly found on the edge of the plot, indicating an obvious edge effect. Thus, sampling should be in the vicinity of the central area to avoid the error caused by the edge effect.

## 5. Conclusions

5.1 Fertilization significantly increases the concentration of MBC and MBN, especially with input of organic materials, and it is increased more in topsoil than sub-topsoil. However, there are few differences of MBC/MBN between topsoil and sub-topsoil and among fertilization treatments.
5.2 Variation of MBC, MBN, and MBC/MBN caused by fertilization were in the following order MBN > MBN > MBC/MBN. Differences in soil properties among fertilization treatments were in the order CK > NPK > NPKM > NPKS (topsoil) and NPK > CK > NPKS > NPKM (sub-topsoil).
5.3 Spatial heterogeneity differed between topsoil and sub-topsoil at the plot-scale. The spatial variability at the plot-scale was NPKM = NPK > NPKS > CK (topsoil) and NPKM = CK > NPK = NPKS (sub-topsoil). Spatial heterogeneity also differed at the fine-scale between topsoil and sub-topsoil: the order was NPK > NPKS > CK = NPKM in topsoil and NPKM > NPK = NPKS > CK in sub-topsoil. Spatial variation of variables followed the order: MBC > MBN > MBC/MBN (topsoil) and MBC = MBN = MBC/MBN (sub-topsoil).
5.4 The “hotspots” of MBC and MBN in the plots were mainly on the plot edges. Sampling should be in the central area of the plot to better represent the situation in plots.

**Fig. 1.**
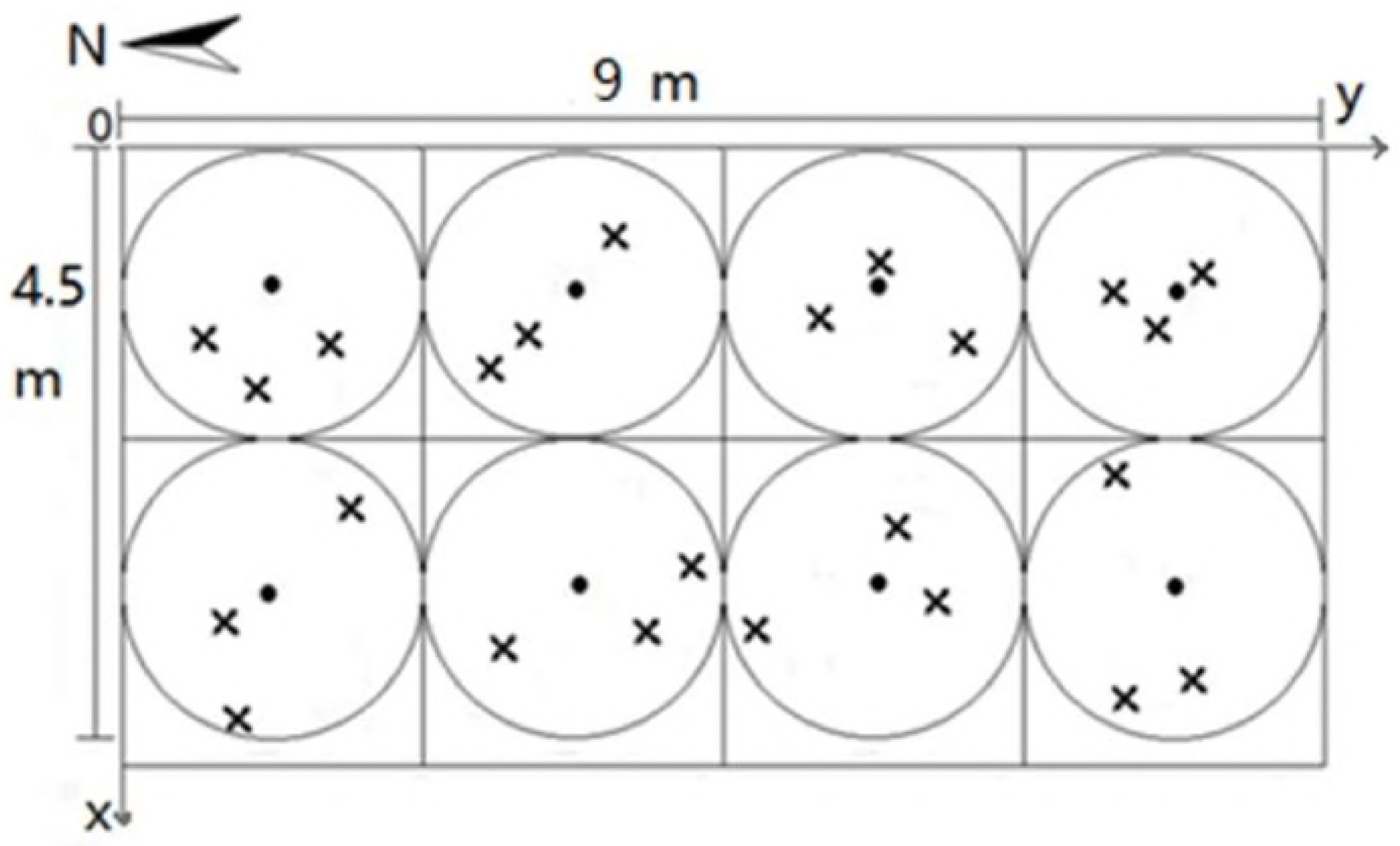
Example of a spatial stratified random sampling design with two transects. Small filled circles represent centroids (n=8), large circles represent the potential sampling area around centroids (radius=1.125m). Xs represent sample locations determined from randomly chosen directions and distances from a centroid. The extent of the interpolation maps based on the inverse distance weighting (IDW) method was determined by the minimum and maximum values at the x and y axes. Therefore, each map had a different extent from the study area (4.5 m × 9 m), which represents the largest extent that each map can attain.

## Acknowledgements

Financial support was received from the National Natural Science Foundation of China (41471248) and the National Key Research and Development Plan (2016YFE0112700, 2016YFD0200301).

